# A Comparison of Antibody-Antigen Complex Sequence-to-Structure Prediction Methods and their Systematic Biases

**DOI:** 10.1101/2024.03.15.585121

**Authors:** Katherine Maia McCoy, Margaret E. Ackerman, Gevorg Grigoryan

## Abstract

The ability to accurately predict antibody-antigen complex structures from their sequences could greatly advance our understanding of the immune system and would aid in the development of novel antibody therapeutics. There have been considerable recent advancements in predicting protein-protein interactions (PPIs) fueled by progress in machine learning (ML). To understand the current state of the field, we compare six representative methods for predicting antibody-antigen complexes from sequence, including two deep learning approaches trained to predict PPIs in general (AlphaFold-Multimer, RoseTTAFold), two composite methods that initially predict antibody and antigen structures separately and dock them (using antibody-mode ClusPro), local refinement in Rosetta (SnugDock) of globally docked poses from ClusPro, and a pipeline combining homology modeling with rigid-body docking informed by ML-based epitope and paratope prediction (AbAdapt). We find that AlphaFold-Multimer outperformed other methods, although the absolute performance leaves considerable room for improvement. AlphaFold-Multimer models of lower-quality display significant structural biases at the level of tertiary motifs (TERMs) towards having fewer structural matches in non-antibody containing structures from the Protein Data Bank (PDB). Specifically, better models exhibit more common PDB-like TERMs at the antibody-antigen interface than worse ones. Importantly, the clear relationship between performance and the commonness of interfacial TERMs suggests that scarcity of interfacial geometry data in the structural database may currently limit application of machine learning to the prediction of antibody-antigen interactions.

## 1. Introduction

When it comes to predicting protein-protein interactions (PPIs), antibody-antigen PPIs are especially important, both to better understand the immune system and for the development of antibody therapeutics. They are a key receptor in the immune system (Nicholson 2016) and genetic variation drives the humoral adaptive immune response in vaccination, infection, and autoimmunity (Mikocziova et al. 2021). Due to this importance, antibodies are actively being researched to understand their correlates of protection for various diseases (Huang et al. 2020) (Plotkin 2023) and ability to neutralize pathogens (Gumah Ali et al. 2020), to utilize antibody indexes to diagnose viral infections (Shamier et al. 2021), to apply them to various types of imaging (Rodriguez et al. 2022) (Im et al. 2019), and to engineer immunotherapeutics for a wide variety of pathologies, such as cancers (Scott et al. 2012) (Zinn et al. 2023; Lucas et al. 2021) and autoimmune diseases (Carter and Rajpal 2022). Research into understanding and engineering antibody binding is largely focused on their Complementary Determining Regions (CDRs), hypervariable regions that predominantly determine their binding repertoires (Peng et al. 2022) (Chiu et al. 2019). Yet, despite much bioinformatic research and progress, predictive modeling of CDR structures and their binding motifs is still a challenge (Bielska-Junior et al.).

To improve our ability to predict these motifs, many antibody-antigen PPI prediction methods have been developed and evaluated on their own, and there have also been some limited head-to-head comparisons of different methods. The rigid body docking programs ZDOCK and ClusPro have been compared head-to-head in their ability to predict complexes from unbound structures, and the local refinement program SnugDock has had its models rescored with ZRANK2 and compared to that reranking (Guest et al. 2021). AlphaFold v2.2.0 has been compared to ZDOCK in its ability to dock AlphaFold-generated antibodies and antigens (Polonsky et al. 2023). AlphaFold-Multimer has also been compared to ClusPro’s ability to dock AlphaFold-Multimer generated antibodies and antigens, although that case only evaluated whether or not models met a highly stringent DOCKQ score cutoff of 0.49 or above (Yin and Pierce 2024). RoseTTAFold has not been evaluated in its capability to predict full antibody-antigen complexes, but has been evaluated in its ability to predict antibodies on their own—in which case it predicts the H3 loop crucial for most binding motifs (Regep et al. 2017) better than ABodyBuilder and comparably to SWISS-MODEL, but the overall 3D structure worse (Liang et al. 2022).

However, the field is still missing a systematic comparison of a diverse array of methods, representative of the variety now available, which can take the sequences of antibodies and antigens as inputs and model their full complex structure as an output. These methods include state-of-the-art general PPI predictors like AlphaFold-Multimer and RoseTTAFold, a number of antibody-specific docking methods, options for local refinement of docked structures, and PPI predictors using machine-learning epitope and paratope prediction to inform their models. Further, it would be most informative to evaluate state-of-the-art options for these diverse methods against each other with less stringent cutoffs than previous work, since methods involving rigid-body docking have shown some success in the past, but are not likely to meet highly stringent DOCKQ scores (which rely heavily on native-like interface recovery) . It would also be useful to examine any systematic biases these various methods may have in terms of structural motifs or amino acid preferences, to identify where the methods go right and where they go wrong. Finally, a good systematic comparison must have a benchmark suitable to both evaluating modern machine learning (ML) methods and antibody-antigen structures in particular, which each require special consideration. Powerful ML methods like AlphaFold-Multimer and RoseTTAFold often have the capability to “memorize” the structures they have trained on (Tsaban et al. 2022)(Rabin et al. 2023) and so require test datasets sufficiently different from those used in training to fairly test for generalization—i.e. to test their ability to predict novel, unsolved antibody-antigen structures. This need is complicated by the particularities of antibody-antigen interactions: antibody binding is almost entirely determined by the CDRs (Peng et al. 2022), while the non-CDR “framework” comprising around 85% of the variable domain of the antibody is highly conserved (Elgert 1998). As a result, filtering benchmark datasets by overall sequence identity to structures trained on is not necessarily appropriate, as it would be in most other PPI prediction evaluations. Moreover, as ML methods train on newer and newer data, benchmark sets will quickly become obsolete. An ideal benchmark would thus be automatically generated with reproduceable standards—such as CDR sequence identity to past training data, sequence identity within the benchmark, resolution of the structures, etc.—so that new benchmarks with precisely the same standards as previous ones may be used to evaluate the improvement of ML methods in particular.

Herein, we address this gap by providing a comparison of six methods for predicting antibody-antigen structures that represent the variety of classes of methods now available, utilizing a benchmark suitable to test ML methods for antibody-antigen prediction with automatically-reproducible standards. We find that AlphaFold-Multimer performs best, RoseTTAFold and AbAdapt perform worst, and the ClusPro and Snugdock methods are in-between. Further, for the winning method AlphaFold-Multimer, we find that inaccurate structural predictions tend to exhibit antibody-antigen interaction geometries that are less frequently found in the PDB relative to more accurate predictions. This may be used to score models based on their structural interaction motifs for a novel confidence measure.

## 2. Results

### 2.1 Model Generation

A benchmark of 57 unique antibody-antigen interactions was generated using a novel method called the Antibody-Antigen Dataset Maker (AADaM), which allows for reproducing benchmark sets of the same standards with different date cutoffs. Structures accepted into the benchmark were required to have below 80% CDR sequence identity with protein-binding antibody structures released past the training cutoff date of every ML method tested, to ensure the benchmark was fair with respect to those methods in that it contained no close similarities to potential training data. Within the benchmark, targets were disallowed from sharing 80% or more sequence identity between heavy chain CDRs, light chain CDRs, or any antigen chains to remove redundancy from the benchmark, for the sake of removing representation bias. The benchmark was filtered for various indications of quality: resolution, method used to determine the structure, gaps in the CDRs or antigen, length of the antigen, etc. (Fig S1, also see Materials & Methods).

The sequences of the benchmark structures were then provided to six methods of antibody-antigen prediction, representative of the different methods currently available. AlphaFold-Multimer and RoseTTAFold are state-of-the-art general PPI-predictors according to the CASP14 and CAMEO benchmarks (Baek et al. 2021), but are not specially trained to account for differences in antibody-antigen interactions. ClusPro is a Fast Fourier Transform-based rigid body docking program with an antibody-specific docking mode, which requires the antibody and antigen to be modeled separately, if the unbound structures are not available. When compared to a similar state-of-the-art rigid body docking program, ZDOCK, it displayed a moderately higher overall success rate (Guest et al. 2021). To predict the partner structures used to dock in ClusPro, we modeled the antigen alone in AlphaFold-Multimer, then tested two approaches for modeling the antibody: using AlphaFold-Multimer as well, or ImmunoBuider, which has been shown to predict H3 loops better than AlphaFold2 (Abanades et al. 2023). Then to test local refinement in SnugDock, the top twenty predictions from ClusPro were used as starting structures. SnugDock is a structural optimizer that has shown success at locally docking antibody-antigen models, requiring a variety of “global” docks, i.e. docks that are widely dispersed over the surface of the antigen in reasonable positions. (Sircar and Gray 2010). (Jeliazkov et al. 2021) It succeeds at locally refining CDR loops more accurately than standard RoseTTADock (Sircar and Gray 2010). Finally, AbAdapt (Davila et al. 2022) is an online pipeline that uses homology modeling to predict the antibody and antigen, then rigid body docks them, informed by machine learning epitope and paratope predictors. It was shown to produce at least one correct structure among its predicted structures 88.4% of the time, but its top-prediction accuracy was not directly evaluated. As it only can predict antibodies including both heavy and light chains, only those benchmark targets (more than half of the test dataset) were tested on it. An overview of the database used, methods tested, rankings of the models, and biases examined is provided (Fig 1).

**Figure 1:**
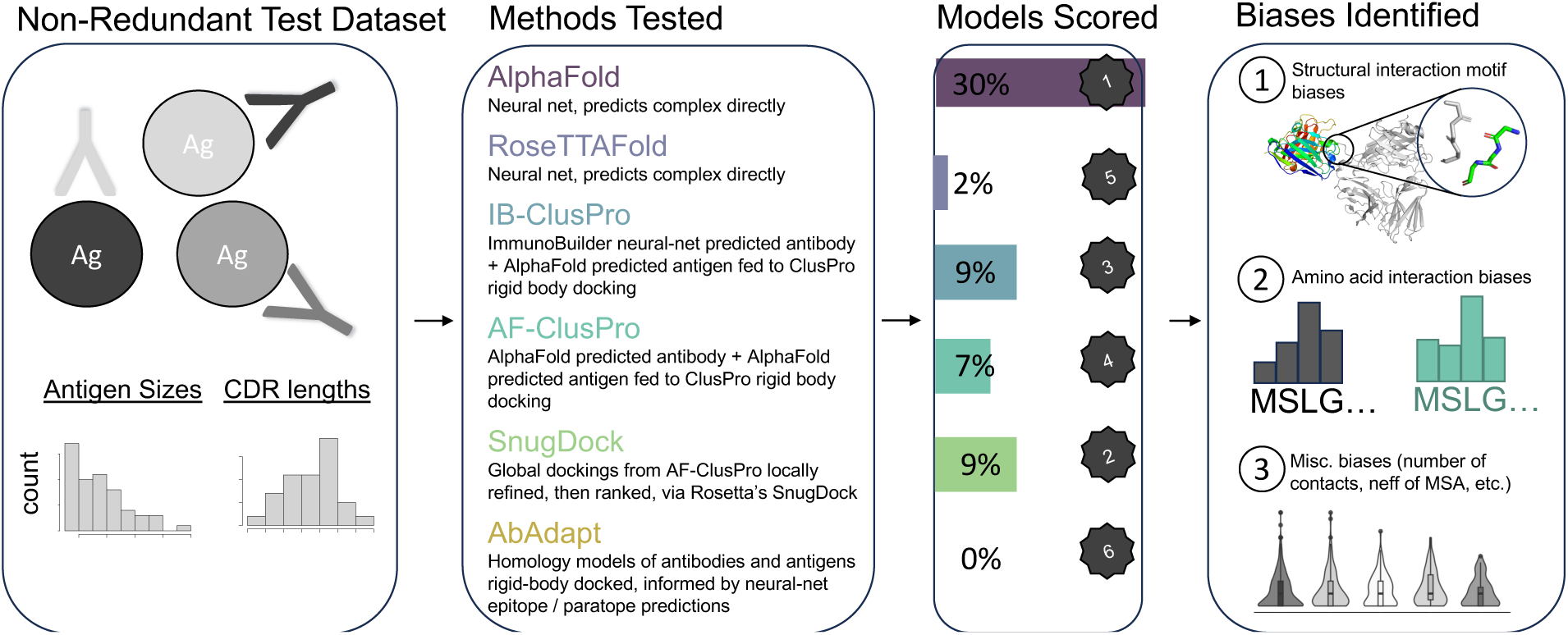
An Overview of How the Antibody-Antigen PPI Methods. *Were Tested and the Biases Examined.* Under “Models Scored”, the overall quality of each ranking according to the percentage of top predictions that were correct by bRD p-value is indicated, with the percentage on the left and ranking on the right.

### 2.1 Model Scoring

The models that each of the six methods produced were then scored in two ways: DOCKQ-score (Basu and Wallner 2016) and biased Random Docking p-value (bRD p-value) (McCoy et al. 2024). DOCKQ score incorporates interface root mean square deviation (RMSD), ligand RMSD (the RMSD of the smaller binding partners upon optimal alignment of the larger binding partners), and number of native interface contacts recovered. A DOCKQ score ≥ 0.23 is commonly used as a cutoff to differentiate correct and incorrect structures (Basu and Wallner 2016), and that cutoff is used here. The bRD p-value measures the likelihood of a binding information-biased random docking to achieve a model with an RMSD as good or better than the model being scored at random. In this case, the random docks were biased to include at least one CDR-antigen contact. It may be interpreted as a straightforward measure of statistical significance like any other p-value. Models with a bRD p-value below 0.01 are classified as “significant” within this paper. These two methods of scoring were used because they are complementary to each other. DOCKQ score is very good at differentiating highly accurate models from each other. However, the ease of achieving a given DOCKQ score varies considerably (> 1000-fold) between different targets, making it difficult to compare between models from benchmark datasets consisting of different structures, which would be highly useful to monitor the improvement of antibody-antigen prediction in the future. DOCKQ is also highly stringent, so that models below the 0.23 cutoff may or may not still be highly significant predictions (McCoy et al. 2024). In contrast, normalizing across different benchmark datasets and identifying highly significant structures classified below a 0.23 DOCKQ threshold are both possible with bRD p-value.

Examining just the top ranked model for each structure in the benchmark, AlphaFold-Multimer outperformed all other methods by both DOCKQ score and bRD p-value: 30% of its top predictions were significant, and 19% of its top predictions were correct (Fig 2a). In contrast, the other general protein structure prediction method RoseTTAFold produced no correct models, and only 2% of its top predictions were significant, only a marginal increase over the 1% o expected to appear significant at random. AbAdapt similarly produced no correct or significant models among its top candidates. In antibody mode, ClusPro docking achieved similar results whether AlphaFold-Multimer (AF-ClusPro) or ImmunoBuilder (IB-ClusPro) was used to model the antibody before docking, with 4-5% of the top models correct, and 7-9% of the top models significant. Performing local refinement with SnugDock on up up to the twenty top structures produced by the AF-ClusPro method actually resulted in slightly worse predictions in terms of top DOCKQ scores, but more significant models overall, when compared to the original AF-ClusPro ensemble and scoring; only 2% of top models were correct but 9% of top models were significant. We also find that incorrect top predictions produced by different methods tend to differ significantly from one another rather than demonstrating convergent erroneous predictions (Table S1), suggesting considerably different innate biases.

**Figure 2:**
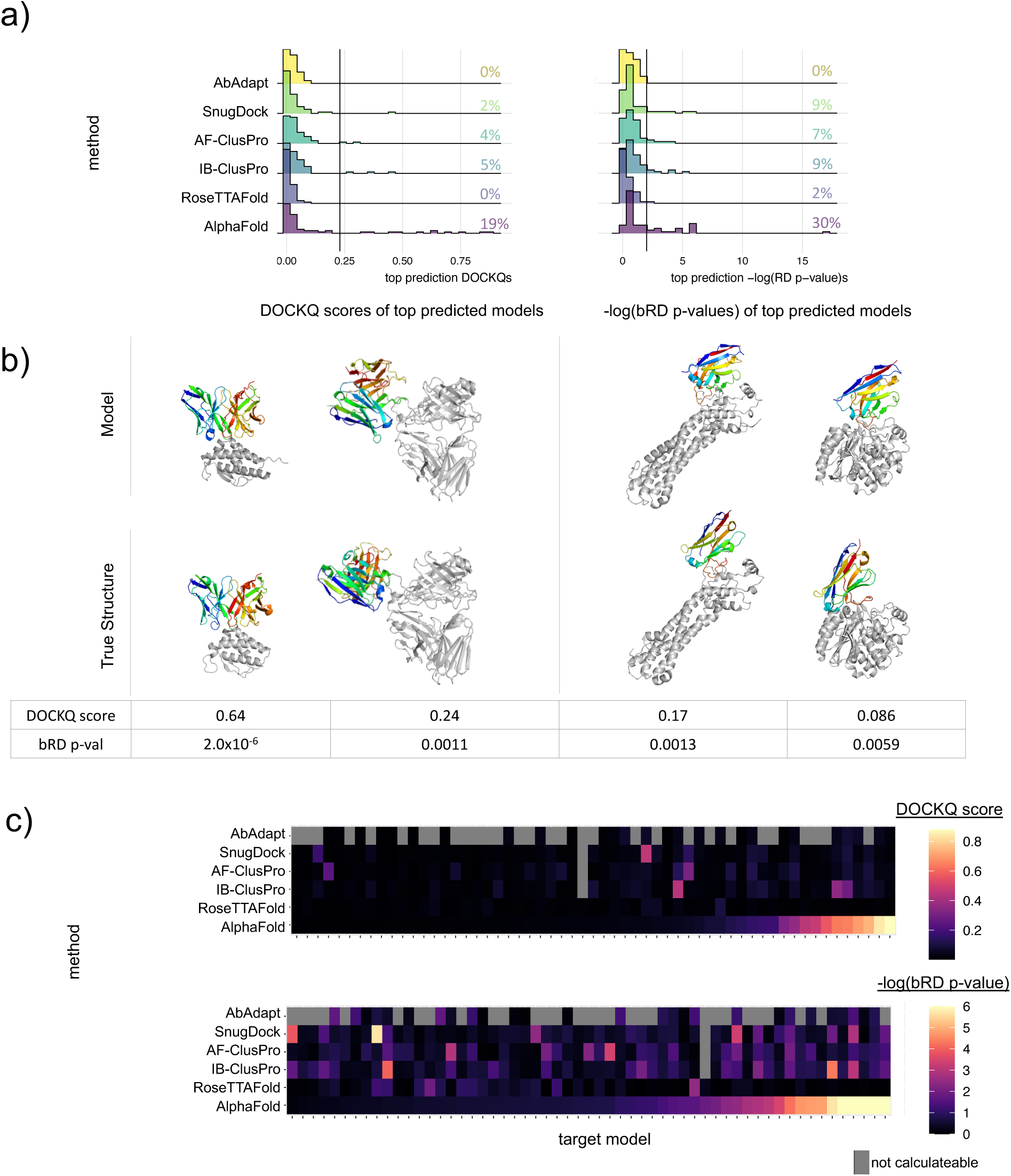
A Head-to-Head Comparison of Top-Ranked Models Shows AlphaFold-Multimer Performs Best at Antibody-Antigen Complex Prediction. a) The top results of each method are compared side-by-side, showing the distributions of their DOCKQ scores and b RD p-values. b) Examples of correct models (DOCKQ scores ≥ 0.23) on the left and approximate models (DOCKQ scores < 0.23, but bRD pvalue < 0.01) on the right are compared to their true structures. Antigens are grey and aligned between true and model structures, while antibodies are rainbow to denote orientation. c) a heatmap lining up each target model in the benchmark set with its score under each method. Grey indicates a model that was unable to be calculated, either because it could not or failed to produce models (in the case of AbAdapt) or the interaction region was trimmed away (ClusPro-based methods), see Materials & Methods for details.

Examples of some models produced in this evaluation, as compared to their true structures, are provided to visualize “correct” models above the 0.23 DOCKQ . Structures that score below that cutoff but that are still highly significant by bRD p-value are also shown, and hereon described as “approximate” models (Fig 2b). Those approximate models have identified the general paratope of the interaction, but have not quite achieved an accurate interface configuration with high enough quality to meet the stringent DOCKQ score threshold. Given the current state of the field, identifying significant but imperfectly -correct structures is worthwhile—it indicates that a given method is picking up genuine signal and succeeding at the “global” docking stage, but not performing adequately at the “local” docking stage. For example, AlphaFold-Multimer produces significant models far more commonly than the 1% that would be expected at random, indicating meaningful success at identifying the general epitope / paratope area, if not the precise conformation of binding.

Comparing the quality of results for the same model between methods (Fig 2c), it’s clear that there is some overlap between methods, but also some orthogonality. The ClusPro-based methods were able to accurately predict some targets that AlphaFold-Multimer did not, indicating a different but useful signal can be detected and used by the rigid body docking than that picked up by the neural net. Interestingly, the top structures with correct DOCKQ scores produced by AF-ClusPro and SnugDock are all predicted incorrectly by AlphaFold, making those methods entirely orthogonal to each other. Similarly, AF-ClusPro, IB-ClusPro, and SnugDock were able to produce significant top structures for some complexes that AlphaFold-Multimer did not.

However, the methods examined do not just produce a single model. Rather, they produce somewhere between 5 to thousands of models, with each model ranked according to its expected quality. AlphaFold-Multimer and RoseTTAFold produce 5 by default, while ClusPro-IB, ClusPro-AF, and AbAdapt produced a variable number of models (usually at least 20 for ClusPro and thousands for AbAdapt). Up to the top 20 models produced by AF-ClusPro were used as starting structure for SnugDock, with each used to produce 50 refined models, for a total of 1000 models per target that were then ranked by Rosetta’s energy score. Looking at just the top five models for each method, AlphaFold-Multimer still outperformed all other methods, producing more significant and more correct models than the others (Fig 3). When considering all five of the models it produced, almost half of the targets presented at least one significant model , and almost a third of the targets presented at least one correct model. As in the previous analysis of top-ranked models, the ClusPro and SnugDock based methods perform worse than AlphaFold-Multimer, but better than AbAdapt and RoseTTAFold. A CDR-biased random docking method would be expected to produce at least one significant-looking model approximately 4.9% of the time at a depth of 5 predictions (see Materials & Methods), so that most of the significant predictions in the top five models of the AlphaFold, IB-ClusPro, AF-ClusPro, and SnugDock methods exceed the success rate expected by chance, whereas the few significant-looking structures produced by AbAdapt and RoseTTAFold may be discounted.

**Figure 3:**
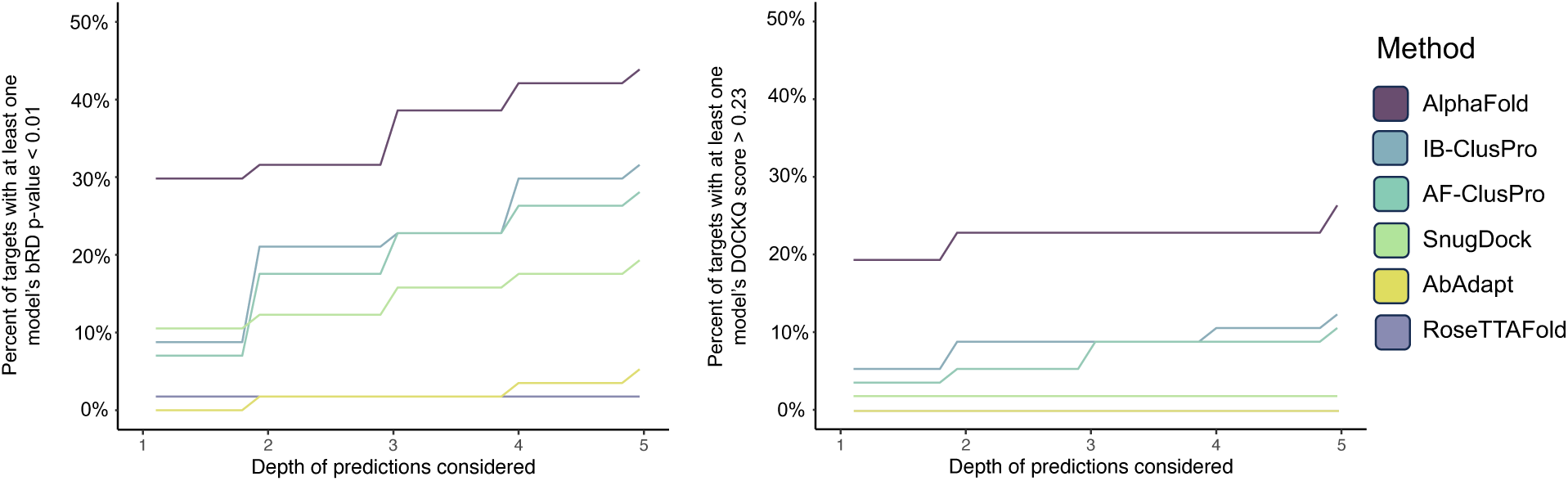
Further Correct and Significant Models are Predicted by Most Methods, But Poorly Ranked. The percent of targets with at least one significant structure (left) or at least one correct structure (right) among the models, at various depths, are shown.

Extending this examination to the top 20 models for each method (Fig S2), IB-ClusPro and AF-ClusPro were able to produce at least one significant model a little over half the time, and SnugDock around a third of the time, which is also more than would be expected by random (18.2%); each of those three methods has a fair number of genuinely significant models that are predicted among the top 20 structures, but simply ranked imperfectly.

### 2.2 Model Biases

After the models were evaluated for accuracy, they were further examined for potential biases. Biases when comparing correct and incorrect models can indicate which factors are important for the success of a model, as well as features that may be used to discriminate good and bad quality models. AlphaFold-Multimer in particular relies on Multiple Sequence Alignments (MSAs), and richer MSAs have been reported to produce better antibody-antigen models (Yin and Pierce 2024). However, as MSAs provide co-evolution information to aid AlphaFold-Multimer in predicting binding, and antigens generally are co-evolving to evade antibodies as well as usually from different species, one would not necessarily expect accuracy in predicting their binding motif to correlate with MSA richness. In line with this, our benchmark dataset showed no significant correlation between the number of effective sequences (Neff) of the MSAs and model quality by DOCKQ score (Fig S3). Paired chain and single chain MSAs were generated automatically by the ColabFold notebook. This difference is likely due to the higher stringency of our benchmark dataset, which had both a higher sequence identity cutoff when comparing to structures AlphaFold trained on, and compared antibodies by CDR sequence identity rather than full variable chain sequence identity.

All methods that produced at least one correct top model were then evaluated for biases between correct and incorrect models in five areas: numbers of residues predicted to be in contact between the antibody and antigen, RMSDs of CDRs, RMSDs of epitopes, structural biases, and sequence-preference biases. No method showed biases in number of contacts predicted between the antibody and antigen; even the rigid body docking methods without flexibility in the CDRs were able to achieve reasonable numbers of contacts (Fig S4).

Increasingly accurate models were also shown to require increasingly lower RMSDs of epitopes and CDRs; however, this difference was more significant for CDR RMSDs, indicating their importance (Fig S5). Most interestingly, there were structural biases revealed by examining the TERtiary Motifs (TERMs) (MacKenzie et al. 2016) making up the interaction motifs of models produced by AlphaFold-Multimer. To extract interaction TERMs, each residue along the antibody that was in contact with the antigen had its nearest antigen residue and both their flanking residues extracted (Fig 4a). Then the interaction-TERM was searched within a 40% sequence identity redundant database of the PDB from which antibodies or antibody-like structures (e.g. TCRpMHC structures) were excluded. To examine if higher quality antibody-antigen structures had more or less common structural motifs, results were split based on the quality of the model the TERM originally came from: incorrect, approximate, acceptable, medium, or high. This analysis was done to evaluate if poor quality models are over-relyiant on common motifs that are not suitable for true antibody-antigen PPIs, which would suggest over-fitting, or if they are simply predicting uncommon geometry when wrong. In order to examine what types of interactions various models are good or bad at predicting in the first place, this analysis was also conducted for the true structures, split by what class of model was produced by the method.

**Figure 4:**
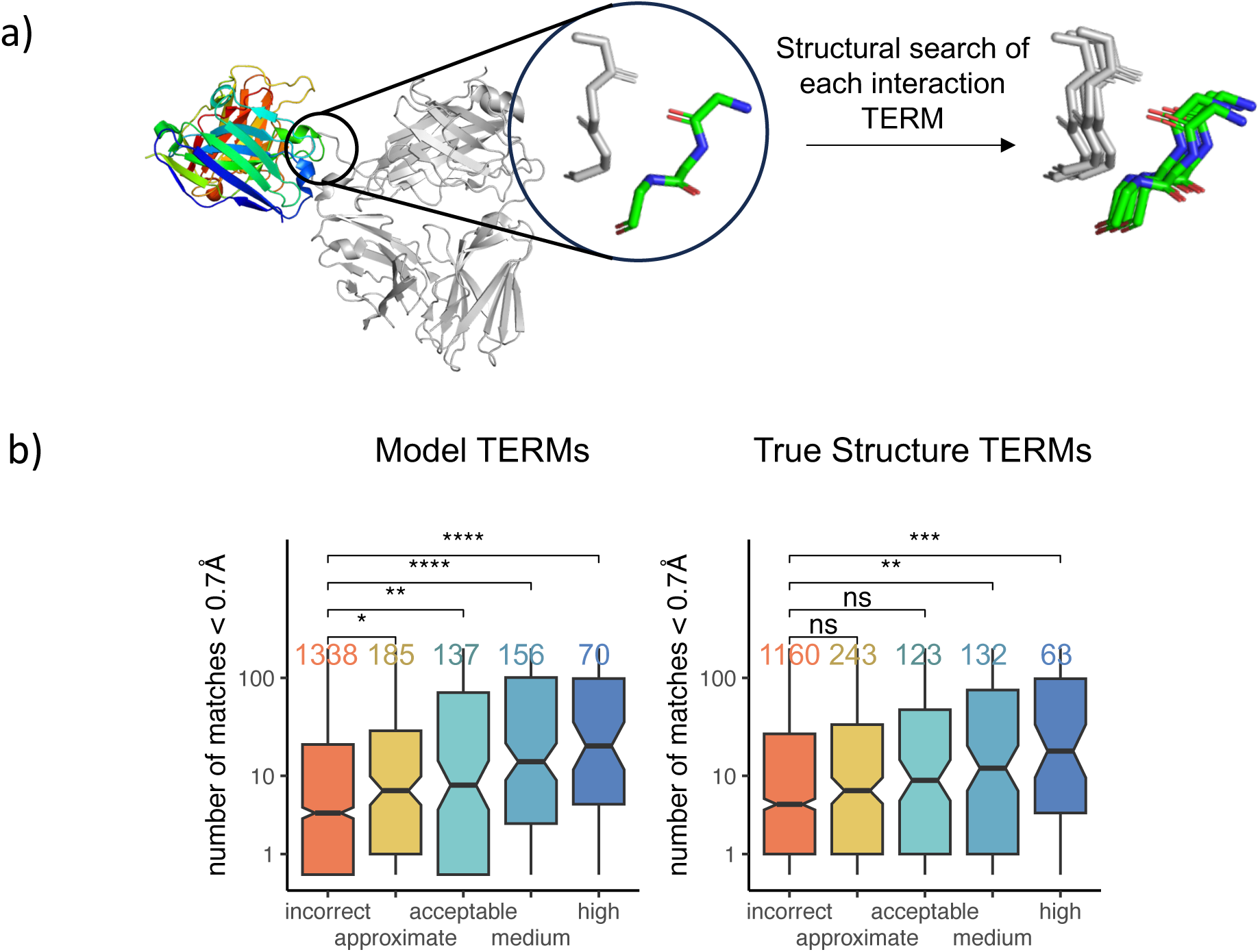
Better Quality AlphaFold-Multimer Models Have More PDB-Like Interaction TERMs. a) an overview of the structural search of a single interaction TERM. The TERM is built from the antibody residue, its closest antigen residue, and all flanking residues. Then it is searched to retrieve close-RMSD matches. b) Interaction TERMs from AlphaFold-Multimer models are split depending on if their original model was incorrect, approximate (significant by bRD p-value, but <0.23 DOCKQ score), acceptable by DOCKQ score, medium by DOCKQ score, or high by DOCKQ score. Then the number of matches below 0.7*Å* is plotted. The notches represent a 90% confidence interval,. This is done for both the TERMs of the model (left) and the TERM of the true structure (right). The number of TERMs evaluated in each category is shown above each box plot in the corresponding color. *\**P *< 0.05, \*\**P *< 0.01, \*\*\**P *< 0.001, \*\*\*\**P *< 0.0001; unpaired two-sided t test*.

AlphaFold-Multimer showed a strong and significant bias towards having more common structural motifs in the interfaces of its more accurate models, and a weaker but still significant bias towards being better at predicting true structures with more common structural motifs in their interfaces (Fig 4b). The trend is stepwise, with the mean number of matches increasing from model classification to model classification; that is, incorrect models have less common motifs than approximate ones, approximate models have less common motifs than acceptable ones. In contrast, none of the other methods showed a similar step-wise trend (Fig S6). The linkage between model quality and common motifs from the PDB indicates that AlphaFold-Multimer is learning, and replicating, interfacial motifs on a TERM-like basis, and has a limited ability to generalize beyond them well.

As large clusters of uncommon structural motifs might be more of an indication that a model is poor than sparsely distributed uncommon structural motifs, this trend was then averaged in local windows to produce a per-model score. Each interaction-TERM and those within 40Å had their closest interfacial match’s RMSD averaged and closest non-interfacial match’s RMSD averaged, then the minimum of each was taken. This minimum was then averaged over the entire interaction motif to create what is hereafter called the CommonInterfacial Motif score (CIM score). Using the CIM score, models can be classified as significant or insignificant with an AUC of 0.725, and correct or incorrect with an AUC of 0.771, which outperforms both average pLDDT and average pLDDT of the interface (Fig 5a). Additionally, CIM score captures very different information than pLDDT, as these scores are not correlated (Fig 5b).

**Figure 5:**
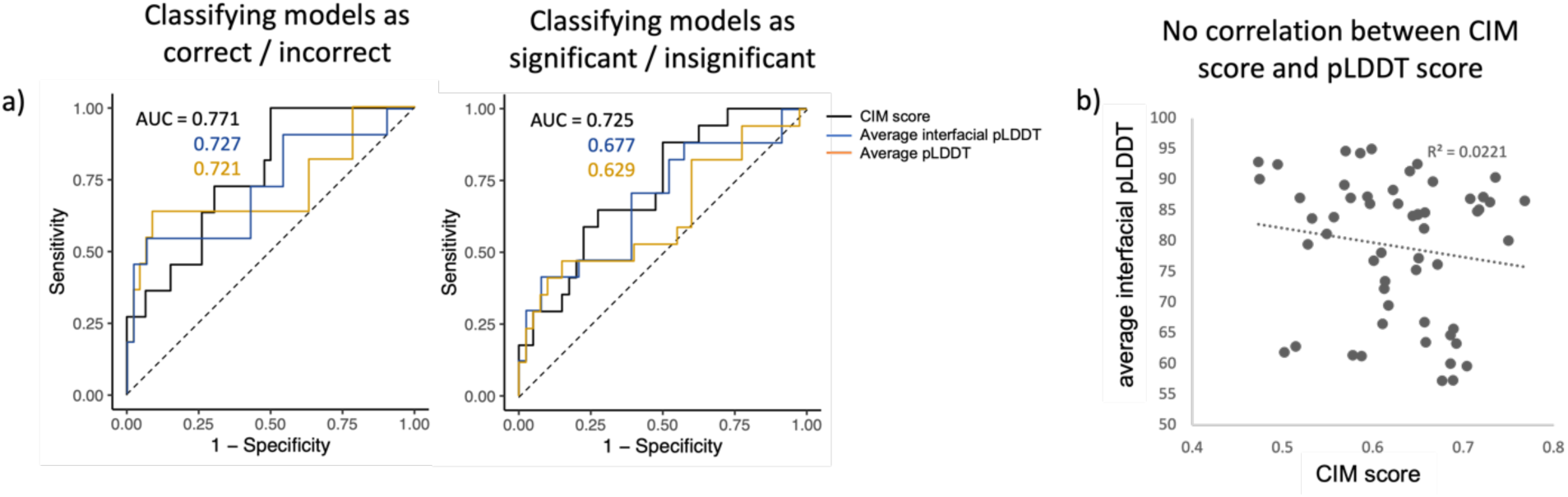
CIM Score Differentiates Good and Poor Structures from Each Other Better than Average Interfacial pLDDT, and Captures Orthogonal Knowledge. a) models are classified as significant / insignificant (left), or correct / incorrect (right), according to CIM score (black), average interfacial pLDDT (blue), or average pLDDT (yellow). b) aggregated best match RMSD of TERMs and average pLDDT do not correlate.

Finally, all methods were evaluated for amino acid biases, in terms of which residues the models preferred to place in contact on either the antibody or antigen side of the interaction motif (Fig S7). In each case, the correct models and incorrect models had their sequence preferences compiled into a distribution, then were compared to the true (background) distribution from the true structures. AlphaFold-Multimer in particular showed a number of significant biases on the antigen side of the interactions in both the correct model ensemble and incorrect model ensemble, despite showing no significant bias on the antibody side for either ensemble (Fig 6). This indicates that although it was trained on general PPI-interactions, it has difficulty with predicting certain residues interacting on the epitope, rather than the paratope. It is especially good at predicting interactions with epitopes high in tyrosine or arginine, or low in methionine, when it succeeds. In contrast, its incorrect models involve epitopes overly high in alanine, glutamine, and methionine, and low in arginine.

**Figure 6:**
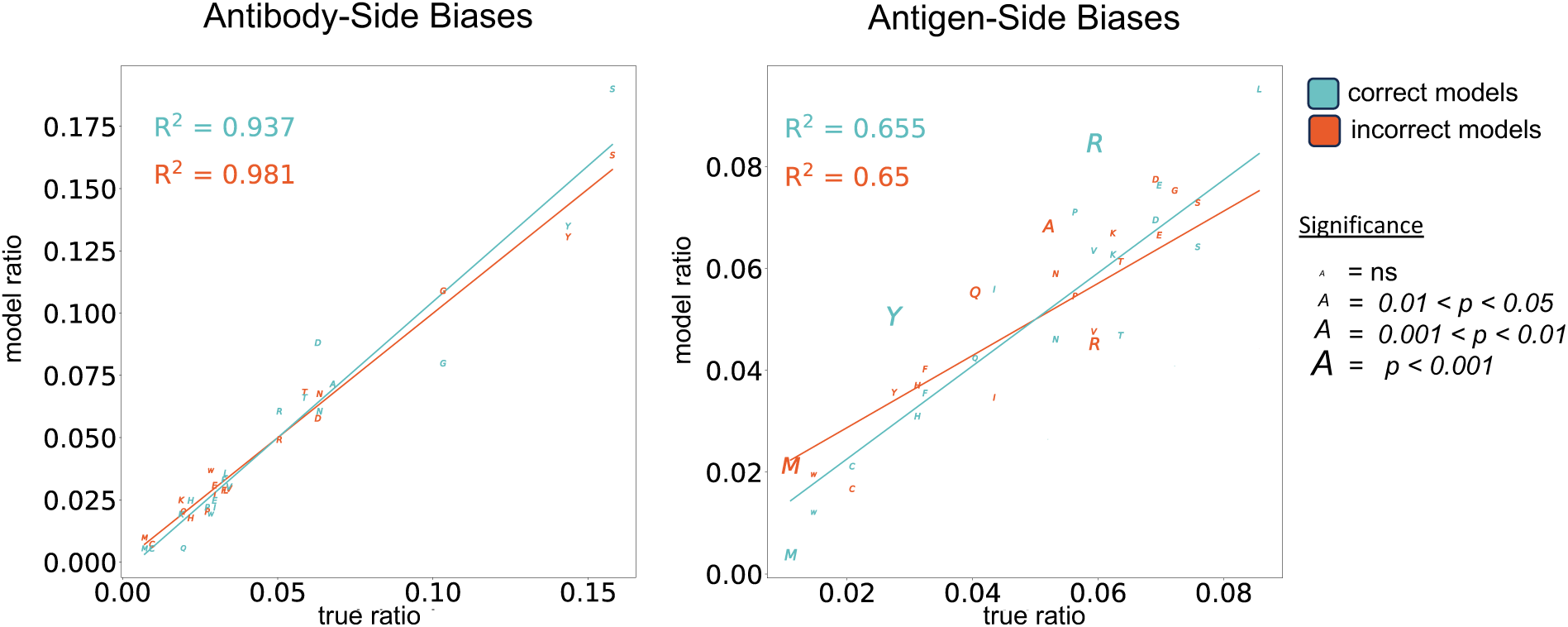
Interaction Residue Biases for Correct and Incorrect AlphaFold-Multimer Models, Compared to the True Distribution. Antibody-side amino acid biases (left) and antigen side amino acid biases (right) are shown. Correct models are in blue, and incorrect ones in red. The size of the letter indicating the amino acid ratio reflects the statistical significance, according to a Fisher’s Exact Test.

## 3. Discussion

On top of being a state-of-the-art general PPI prediction method, our analysis shows that AlphaFold-Multimer is able to out-perform a variety of other antibody-antigen specific PPI prediction methods. This is despite the fact that its top ranked prediction is only correct by DOCKQ score 19% of the time or significant by bRD p-value 30% of the time, in contrast to its to success at predicting general heteromeric interfaces (top prediction correct by DOCKQ score in 70% of cases) (Evans et al. 2022). RoseTTAFold’s inability to predict any of the benchmark dataset correctly is also notable, although this evaluation was performed on RoseTTAFold and not RoseTTAFold2, which may have improved performance on antibody-antigen structures—as it has improved performance when compared to RoseTTAFold overall (Baek et al. 2023). But as the field of machine learning is improving rapidly at antibody modeling, with many advancements like AlphaFold-Mutimer (Evans et al. 2022) and ABodyBuilder (Abanades et al. 2023) developed in only the past few years, it’s likely that ML-based antibody-antigen prediction will improve even further in the near future. Open source platforms like Uni-Fold (Li et al. 2022) offer the opportunity to train attention-based PPI prediction methods from scratch, e.g. on training data biased towards antibody-antigen complexes, or modified for transfer learning. Diffusion-based methods such as Chroma (Ingraham et al. 2023) and DiffDock-PP (Ketata et al. 2023) offer a novel architecture for predicting protein complexes. ML methods performing docking such as DiffDock-PP may be useful for predicting targets that AlphaFold-Multimer currently fails on, as suggested by our discovery of ClusPro’s orthogonal ability to succeed on some targets where AlphaFold-Multimer failed using even rigid-body docking. AlphaFold-Multimer has also very recently been shown to improve at its antibody-antigen predictions with multi-seed runs, on a small benchmark dataset (Yin and Pierce 2024).

As the field develops these improved methods for antibody-antigen PPI prediction, our research suggests that analyzing the interfaces of models on a structural TERM level and a residue-preference level would be particularly useful for monitoring how biased a method trained on general PPI interactions is. Not only did models of good-quality appear more like non-antibody structures from the Protein Data Bank (PDB) the higher their quality was, but this trend can be used per-model to produce a TERM-based quality score that differentiates good and bad antibody-antigen models from each other even better than pLDDT, AlphaFold-Multimer’s own metric of how accurate it expects a model that it produced to be. Moreover, as a structure’s TERM-based score does not correlate with its pLDDT, they are capturing largely orthogonal information; both could be combined to better analyze and rank output models.

Additionally, when considering all five models for each complex that Alphafold-Multimer produced in our analysis, almost half of the targets had at least one significant model, and almost a third of the targets had at least one correct model. This indicates further opportunity for improving AlphaFold-Multimer in particular, both via local refinement of the CDR conformation (e.g. improving those approximate structures that have almost recovered the correct binding motif), and in retraining the ranking for antibody-antigen models.

Finally, AADaM should prove useful for evaluating any future antibody-antigen prediction methods fairly and comparably to past evaluations, providing a consistent report on the state of the field. Its ability to create benchmark datasets with different structures yet the same metrics, in terms of sequence identity cutoffs to previously-trained upon structures, sequence identity cutoffs within the benchmark structures, resolution, and methods, are especially suitable for testing ML methods. There is one other antibody-antigen dataset maker that was recently released (Zhao 2024) and may also be automatically regenerated, titled the ABAG-docking dataset, but AADaM contrasts with it in several different ways. AADaM allows for sequence identity filtering to structures before its cutoff date in order to prevent the test set from containing structures too similar to training data. AADaM filters by CDR sequence identity on the antibody side, while the ABAG-docking dataset filters by overall sequence identity.

AADaM does not collect bound / unbound versions of the individual binding partners, while the ABAG-docking dataset does. Thus AADaM is better designed for fairly evaluating machine learning-based methods, while the ABAG-docking dataset is better designed for evaluating docking methods that use true structures to dock rather than models. AADaM may also be uniquely useful to develop complementary training datasets to its testing datasets, by adjusting the date cutoffs to only allow structures before those of a given benchmark dataset, and thus aid in developing novel ML antibody-antigen prediction methods.

## 4. Materials & Methods

### 4.1 The Antibody-Antigen Dataset Maker (AADaM) Benchmark

Antibodies from the Structrual Antibody Database (SAbDab) (Dunbar et al. 2014; Schneider et al. 2022) were downloaded on Jaunary 17th 2023 and provided to AADaM. Single-Chain Fragment Variable (scFv) antibodies were excluded, as many methods predict the heavy and light chains separately, but DOCKQ score cannot pair multiple chains to a single chain. Only antibody-antigen complexes released after March 14th, 2022 were allowed, as the version of AlphaFold-Multimer tested used templates up to March 13th 2022, and was the most recent cutoff date among other ML methods (RoseTTAFold, AbAdapt). AADaM was used to create a benchmark of structures and their corresponding FASTA files automatically filtered according to the following standards: Structures with >80% CDR sequence identity to antibodies in complex with proteins or peptides released before the Marth 14th cutoff date were disallowed.

Structures from methods besides X-ray crystallography or cryo-EM were disallowed. Structures with a resolution ≥3.5Å were disallowed. Within the dataset, structures were required to have 80% or less sequence identity between heavy chain CDRs, light chain CDRs, and any antigen to antigen chain comparison by global sequence alignment. When structures needed to be discarded to meet these criteria, the one with fewest missing residues within the CDRs and antigens was given preference to remain within the dataset. If both shared the same number of missing residues, the structure with the shorter antigen sequence was selected. Sequences with non-standard residues on their ends had those residues stripped, and sequences with non-standard residues in the middle were disallowed. Only the variable regions of the antibody sequence were saved in order to simplify antibody modeling for all methods.

Finally, the database was parsed by hand to remove structures with an antigen size of 1500 residues or more, as RoseTTAFold is unable to predict structures that large and they are more computationally expensive to all methods.

AADaM is available on GitHub (https://github.com/kaymccoy/AADaM).

### 4.2 AlphaFold-Multimer (AlphaFold) Modeling

AlphaFold-Multimer models were generated using the ColabFold notebook (Mirdita et al. 2022) with the MMseqs2 (UniRef+Environmental) MSA mode, the templates option, 5 models generated, 3 rounds of recycling, and the AMBER (Hornak et al. 2006) relaxation option. Its most recent update was titled 10Oct2022; AlphaFold-Multimer had been trained on Uniref 2022_02 and searched for templates within the PDB/PDB70 220313. For brevity, this method is referred to in figures as “AlphaFold”.

### 4.3 RoseTTAFold Modeling

RoseTTAFold models were generated using the Robetta webserver (Hiranuma et al. 2020; Baek et al. 2021) in February 2023. The webserver was provided MSAs with only those chains from the antibody paired or those chains from the antigen paired (the rest of the sequence left as gaps). Pairing of the antibody chains and antigen chains was done according to the MSAs produced by ColabFold when predicting the antibody and antigen structures separately (see 4.5). This choice was made according to RoseTTAFold’s documentation and support, which advised doing so for cases in which a complex involved partners that did not mutually co-evolve to bind each other. Using the full paired MSAs from the AF-M runs was also attempted for comparison, but most RoseTTAFold runs failed to complete in that case.

### 4.4 ImmunoBuilder Antibody Modeling

ImmunoBuilder (Abanades et al. 2023) was used to model the antibodies in the AADAM benchmark using ABodyBuilder2 for those antibodies with both heavy and light chains, and NanoBodyBuilder2 for the nanobodies. These were then provided to ClusPro for rigid body docking (see 4.6).

### 4.4 AlphaFold-Multimer Individual Antibody or Antigen Modeling

AlphaFold-Multimer was also used to separately model the antibodies and their antigens from the 57 structures comprising the AADAM-generated benchmark, using the same ColabFold web server and options as when modeling the full complexes (previously described in 4.2).

Additionally, the ends of each chain on the models were trimmed until a residue with a pLDDT ≥90 was reached. Docking with untrimmed models was also attempted, but it caused ClusPro to crash on 7 out of 57 attempts, whereas trimming resulted in only one antigen having its interface-region trimmed away so that it was unable to be scored, making the overall results slightly better (Fig S8). The final method compared in the main text used the trimmed antibody and antigen models for docking. Antibodies were also ImMunoGeneTics (IMGT) numbered using Antigen receptor Numbering And Receptor ClassificatIon (ANARCI) (Dunbar and Deane 2016).

### 4.6 ClusPro Antibody-Mode Docking

ClusPro docking was performed using the webserver, with the antibody-mode and non-CDR masking options checked (Brenke et al. 2012; Kozakov et al. 2013; Kozakov et al. 2017; Vajda et al. 2017; Desta et al. 2020). Either ImmunoBuilder-predicted antibodies were docked with AlphaFold-Multimer predicted antigens, referred to as “IB-ClusPro” in figures for brevity, or AlphaFold-Multimer predicted antibodies and antigens were docked, referred to as “AF-ClusPro”.

### 4.7 SnugDock Local Docking

Up to the top 20 structures from the AF-ClusPro method were provided to SnugDock (Sircar and Gray 2010; Jeliazkov et al. 2021). Each was prepacked with Rosetta’s default prepacking options, then used to generate 50 refined structures, for up to 1000 structures per model. The snugdock flags used were: -nstruct 50, -dock_pert 3 8, -detect_disulf false, -ex1, -ex2aro, - input_ab_scheme IMGT, -docking_local_refine true, and -spin. This method is referred to as “SnugDock” in figures for brevity.

### 4.8 AbAdapt Modeling

AbAdapt Models were predicted using its online pipeline (Davila et al. 2022). As AbAdapt may only predict antibodies with both heavy and light chains, those 31 of the 57 structures in the benchmark were tested, and percentages were calculated out of 31 rather than 57. Out of those 31 pipeline submissions, 4 failed with errors indicating the antibody or antigen had failed to model, and 27 succeeded in producing models.

### 4.9 bRD p-value scoring

bRD p-values (McCoy et al. 2024) were used to evaluate the significance of receptor + ligand RMSD for every model examined. An ensemble of 500,000 random docks were used per score, and the random docks were biased to require at least one CDR residue to be in contact.

A random docking process would be expected to achieve a model with a significance of 0.01 or below 1% of the time, so that if X random models are considered, at least one of those models would be expected to appear “significant” at a rate of 1-(0.99X). This expected value offers a reference random expectation, which provides useful context to interpreting ensembles bRD p-values.

### 4.10 CDR Centroid Distance Calculation

All pairs of incorrect top models, between different methods as well as the true structure, were compared. Each model was aligned to the true structure by the antigen. Then the center of mass of the CDRs, according to IMGT numbering, was determined, and the distance between centroids was calculated.

### 4.11 MSA N_eff_ Bias Analysis

The number of effective sequences for MSAs (Neff ) was calculated using a sequence cutoff of 80%, as previously performed in evaluating AlphaFold (Evans et al. 2022). Neff was taken for the paired MSAs and averaged for the single chain MSAs, then both average-single and paired Neff was correlated with DOCKQ score.

### 4.11 Contact Number Bias Analysis

To count the number of contacts predicted between the antibody and antigen in each model examined (the top predicted models of AlphaFold-Multimer, AF-ClusPro, AB-ClusPro, and SnugDock) contact was determined according to contact degree (Zhou et al. 2020). Any residue with a ≥0.001 percent overlap between rotamers was counted as a contact.

### 4.12 TERtiary Motif Bias Analysis (CIM score)

The interaction TERMs were constructed by iterating over contacts as described in 4.11, then constructing a TERM out of each antibody residue and only its closest antigen residue contact, plus their flanking residues. Each TERM was then searched within a 40% sequence identity cutoff redundant version of the PDB downloaded on June 18th 2021, using the MASTER v 2. search algorithm (Zhou and Grigoryan 2020), as implemented in the GrigoryanLab/Mosaist Github Repository, after removing any antibody or antibody-like structures (such as T Cell Receptor complexes). Antibody or antibody-like structures were determined by whether or not they were recognized and numbered by ANARCI (Dunbar and Deane 2016).

The distance cutoff used for finding structural matches (≤0.7Å) was determined by testing at a range of possible cutoffs, then choosing the one with best discrimination of AlphaFold-Multimer structures. However, discrimination was largely robust to this choice (Fig S9). For time, a maximum number of matches was applied. Up to 100 non-interfacial matches (matches between segments on the same chain) and 100 interfacial matches (matches between segments on different chains) were accepted. Differences between interfacial matches and non-interfacial matches were compared, but ultimately combined as they both showed the same trend (Fig S10,S11).

To create a CIM score for each structure, each interaction TERM and those within 40Å had their closest interfacial matches’ RMSD averaged and closest non-interfacial match’s RMSD averaged, then the minimum of each was taken. This was then averaged per structure.

### 4.13 Sequence Bias Analysis

The residues determined to be in contact (see 4.11) were recorded, and categorized based on whether or not they came from a correct (DOCKQ ≥ 0.23) or incorrect model. The sequence preferences of correct and incorrect models were then compared to those of the true structures from the benchmark to look for biases. To check if biases were significant or not, a Fisher’s exact test comparing the preference for the amino acid vs the preference for all other amino acids, between the model category (either correct or incorrect) and the true structure, was calculated.

## Acknowledgments

This work was supported in part by the National Institutes of Health grant R01GM132117, NIAID P01AI162242, and the Dartmouth International Vaccine Initiative.

