## Supplemental Material for "A Comparison of Antibody-Antigen Complex Sequence-to-Structure Prediction Methods and their Systematic Biases"

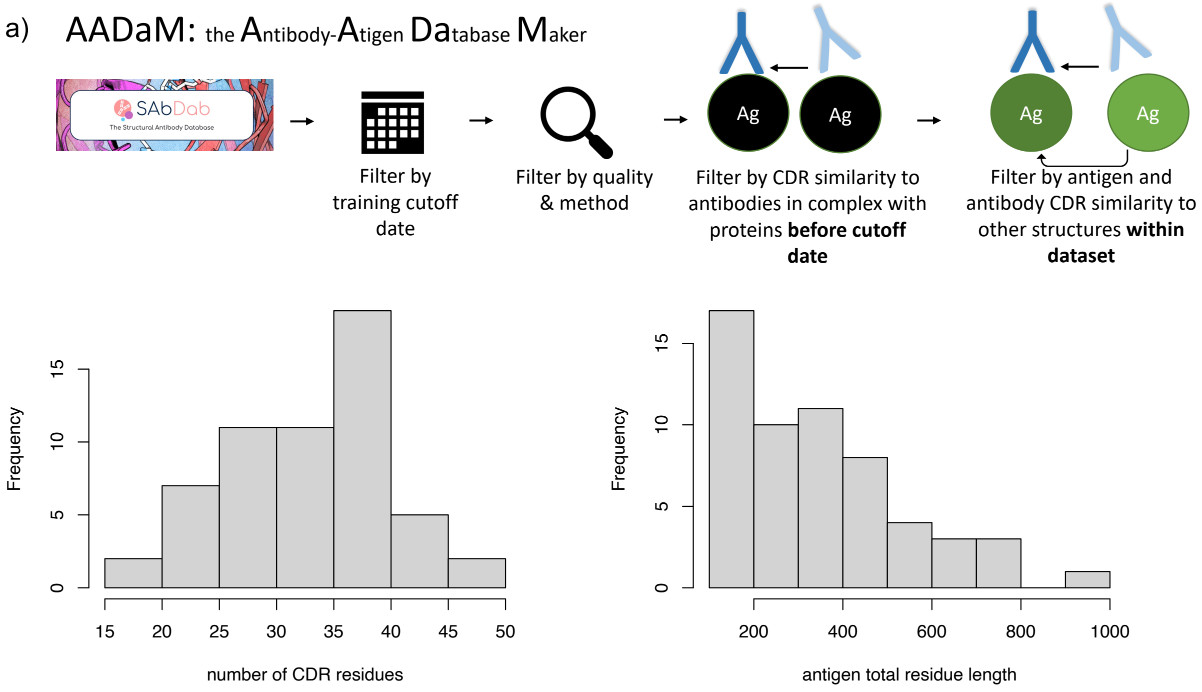


*Figure S1: An Overview of the benchmark produced by AADaM.* a) an overview of AADaM's process. b) distributions of the numbers of CDR residues in the antibodies, and number of total residues on the antigens

| | True Structure | AlphaFold | RoseTTAFold | IB-ClusPro | AF-ClusPro | SnugDock | AbAdapt |
|---|---|---|---|---|---|---|---|
| True Structure | 0 | 42 | 58 | 41 | 43 | 43 | 54 |
| AlphaFold | 42 | 0 | 39 | 31 | 28 | 31 | 31 |
| RoseTTAFold | 58 | 39 | 0 | 40 | 45 | 43 | 42 |
| IB-ClusPro | 41 | 31 | 40 | 0 | 25 | 31 | 34 |
| AF-ClusPro | 43 | 28 | 45 | 25 | 0 | 29 | 26 |
| SnugDock | 43 | 31 | 43 | 31 | 29 | 0 | 26 |
| AbAdapt | 54 | 31 | 42 | 34 | 26 | 26 | 0 |

*Table S1: Incorrect Top Models are Diverse in Their Binding Locations. Shown are median distances between all incorrect predictions by pairs of methods. A distance between two predictions is defined as the CDR centroid distance after superimposing the predictions by the antigen*

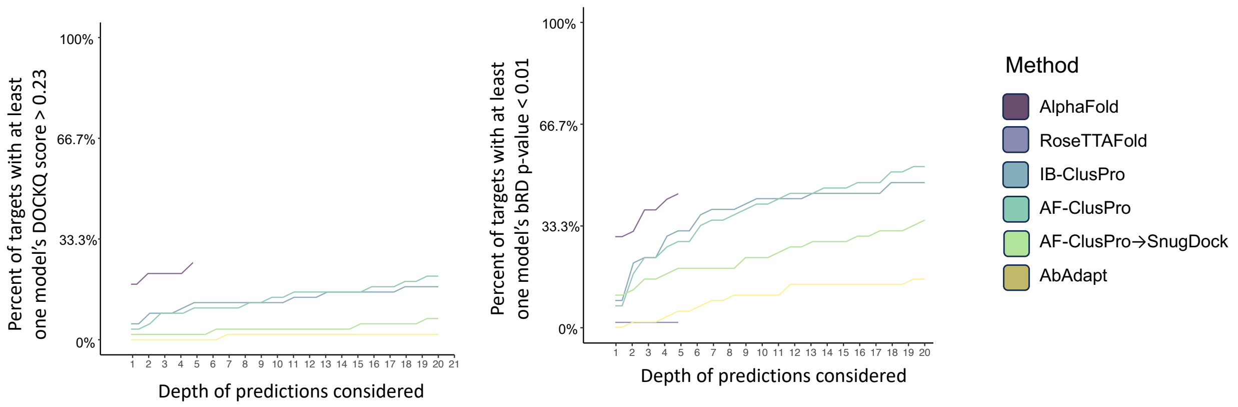


*Figure S2: Cumulative precents of targets with at least one significant (left) or correct (right) structure, at various depth of predictions considered. AlphaFold-Multimer and RoseTTAFold results only included five structures, and so end at that point.*

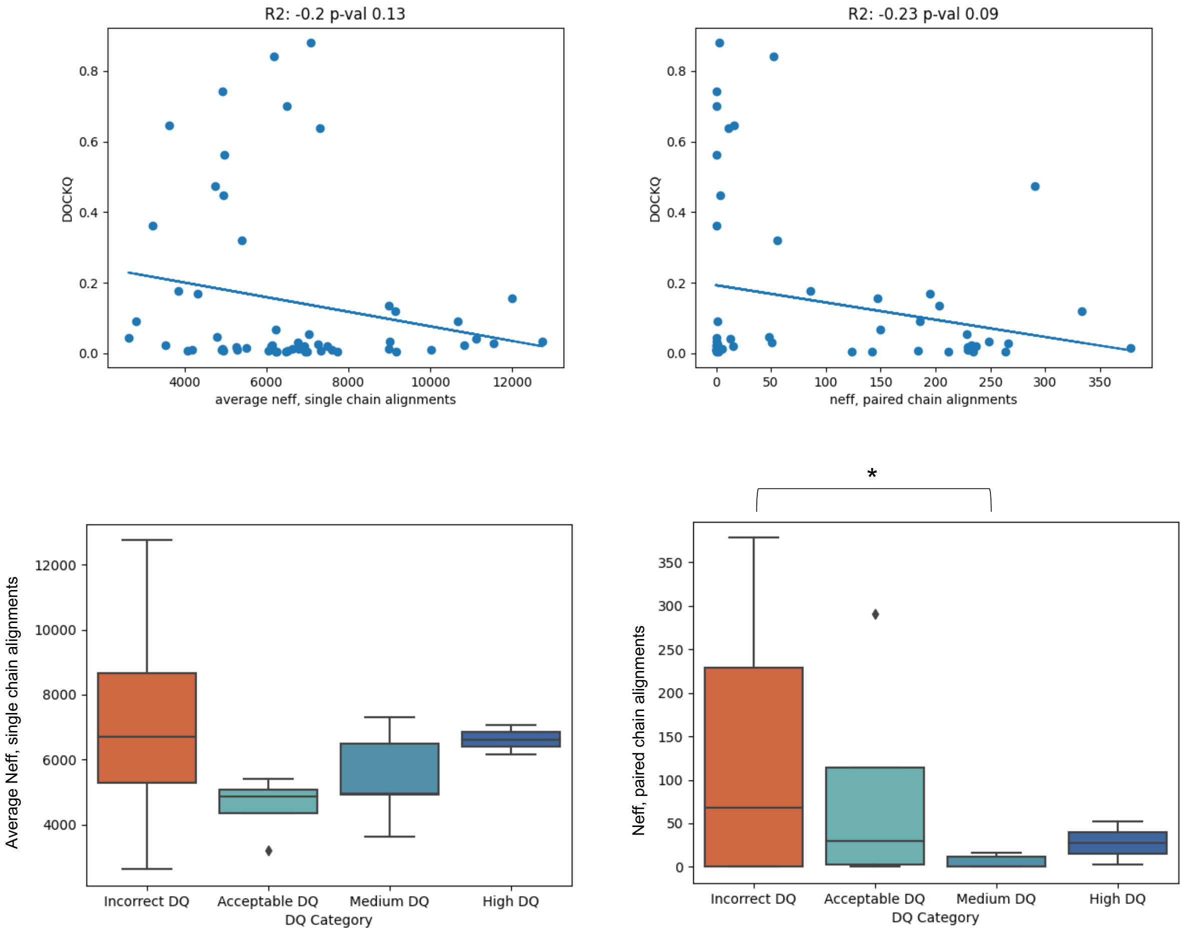


*Figure S3: Analysis of Model Quality Compared to Number of Effective Sequences in its MSA.* $N_{eff}$ of paired (left) and average $N_{eff}$ of single (right) alignments is not strongly correlated with DOCKQ score. Top shows a scatter plot correlation, while bottom splits distributions by DOCKQ score category (incorrect, acceptable, medium, and high).

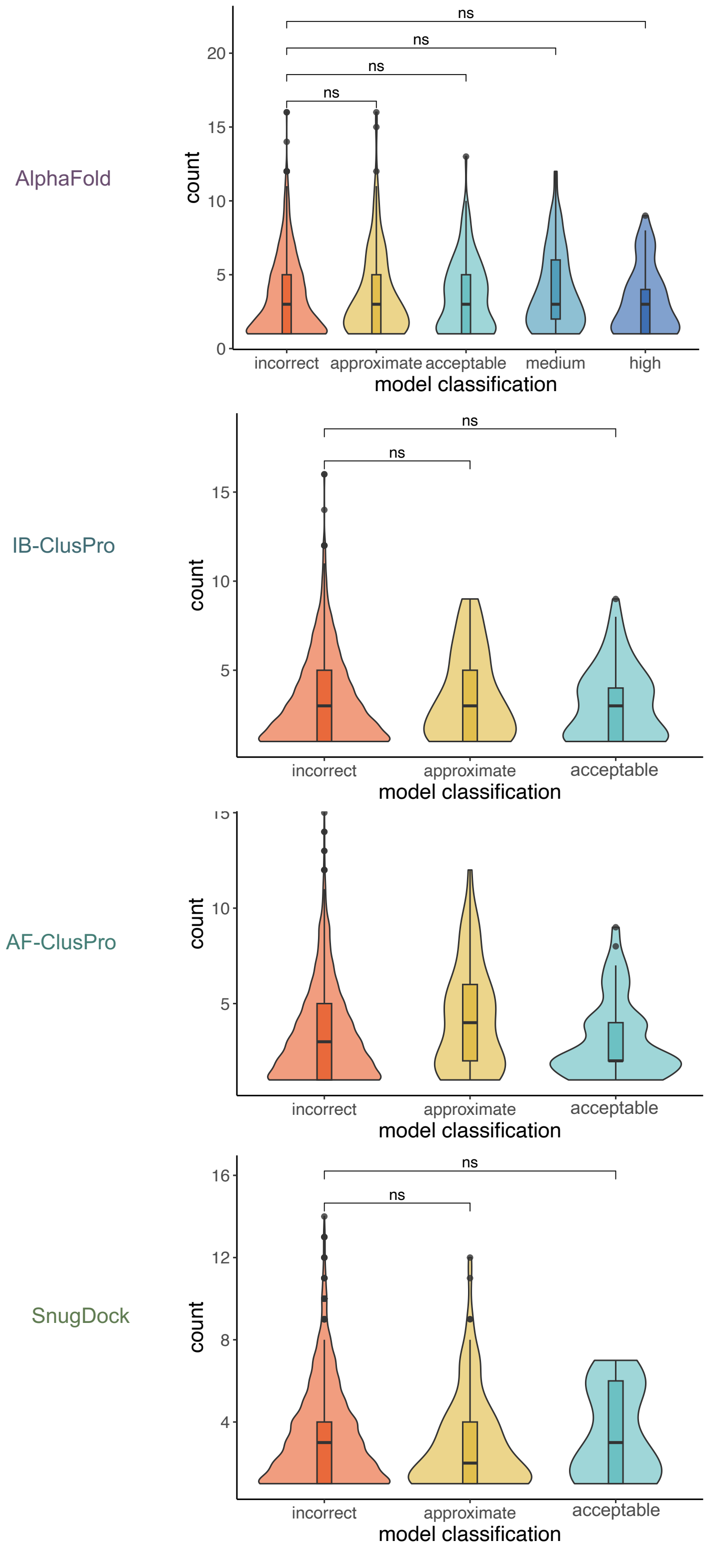


*Figure S4: Analysis of Model Quality Compared to Number of Interactions.* Model quality compared to the number of unique pairs of interactions between the antibody and antigen are shown.

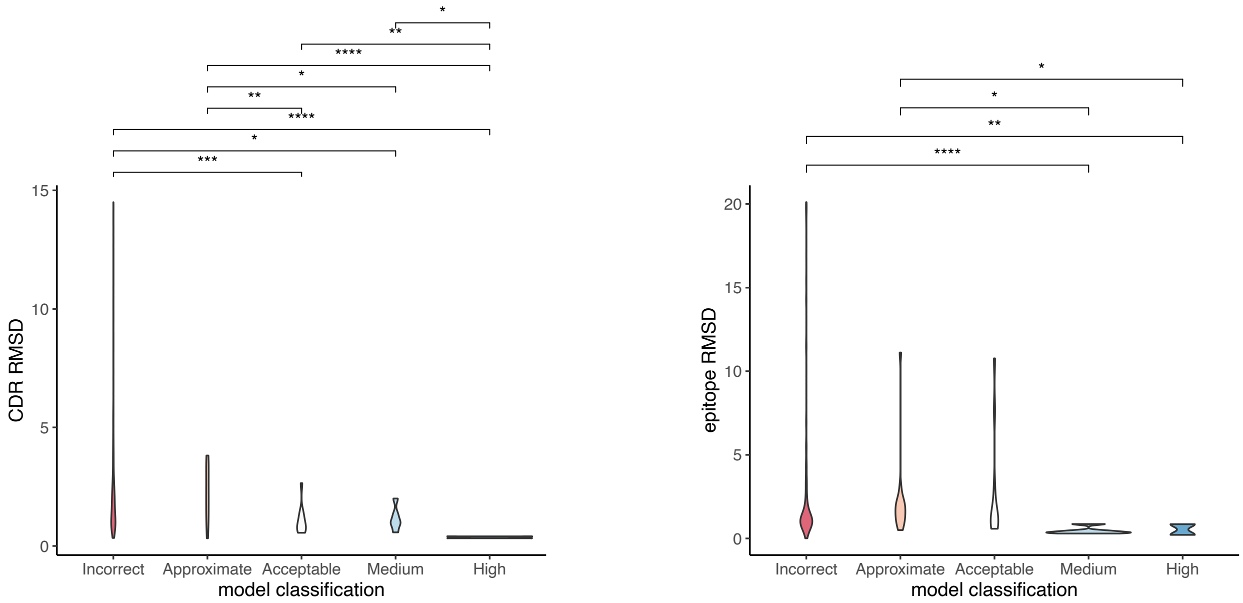


*Figure S5: CDR RMSD Accuracy Is Significantly Important in Producing a High Quality Antibody-Antigen Model*. Model quality compared to the CDR RMSDs (left) and epitope RMSDs (right) is shown.

b)

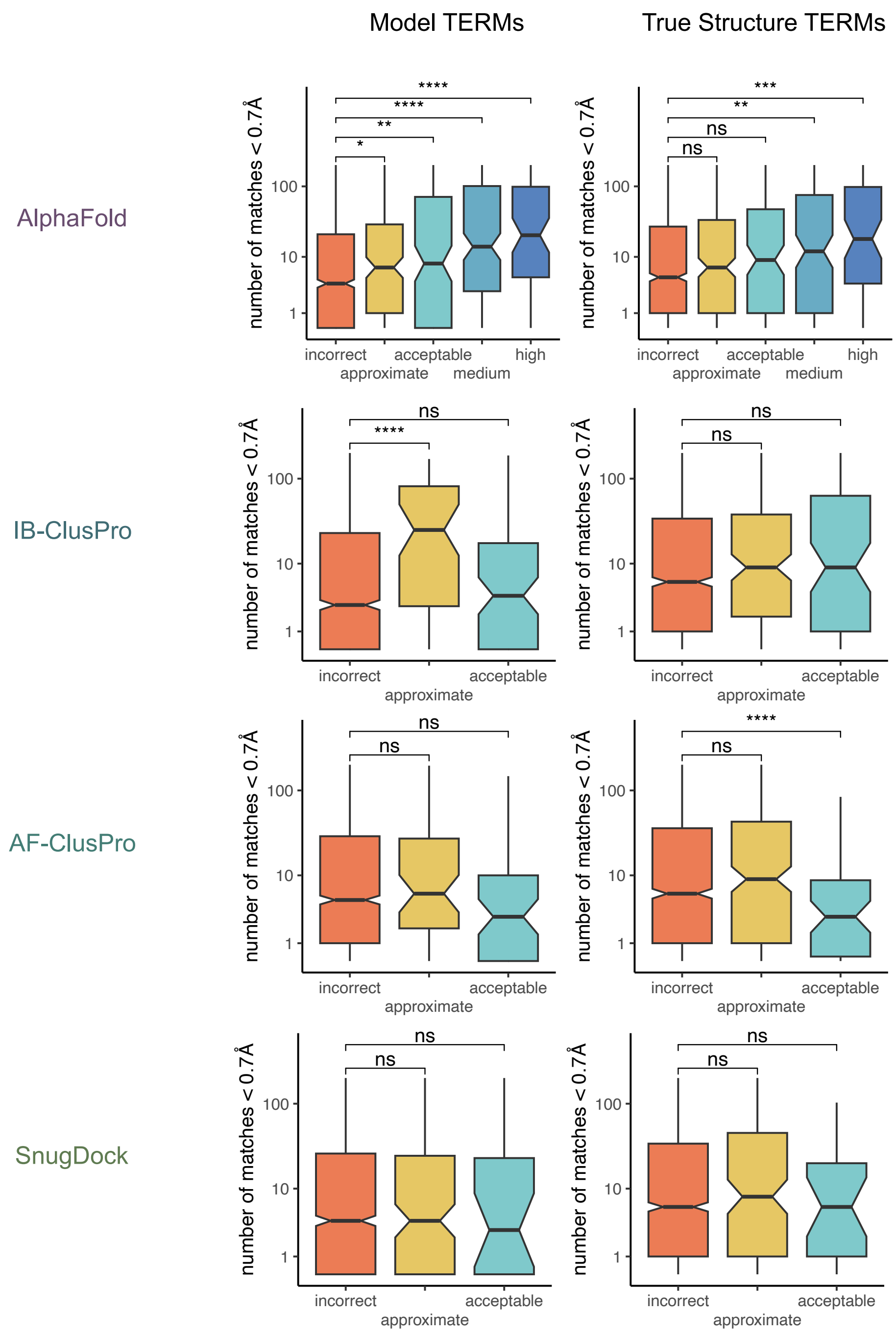


*Figure S6: Structural Biases for Other Methods Producing At Least One Correct Top Structure*. Interaction TERMs from each method's models are split depending on if their original model was incorrect, approximate (meaning significant, but not correct), acceptable, medium, or high. Then the number of matches below 0.7*Å* is plotted. The notches represent a 90% confidence interval, and the colored regions represent the Interquartile Range. This is done for both the TERMs of the model (left) and the TERM of the true structure (right). Significance between incorrect and other ranks of models are indicated with asterisks (* = < 0.05, ** = < 0.01, *** = < 0.001, **** = < 0.0001).

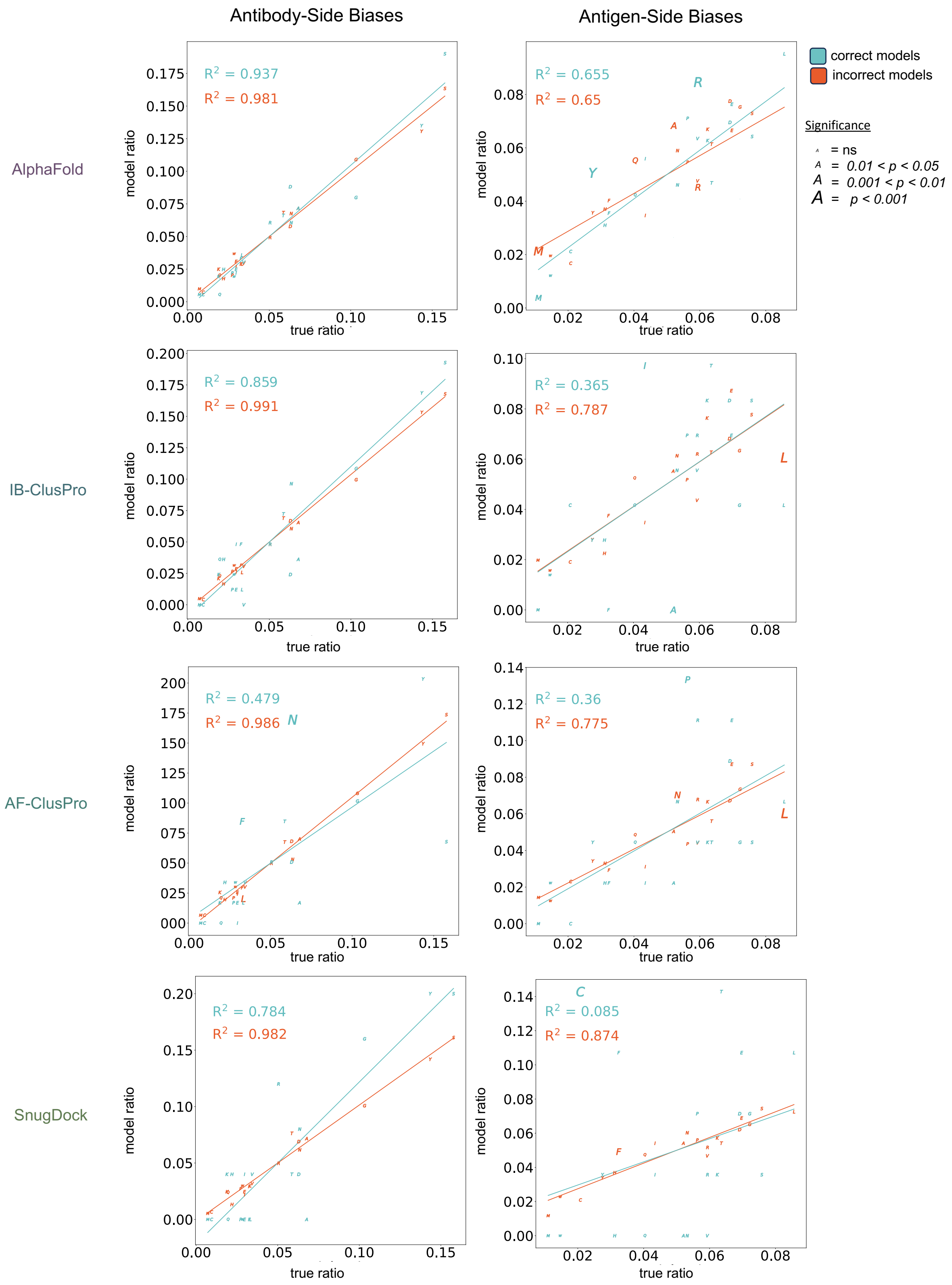


*Figure S7: Sequence Biases for Correct and Incorrect Models from Various Methods*. Antibody-side (left) and antigen-side (right) sequence biases are shown, with each letter representing the amino acid, and its position representing the true ratio compared ot the model ratio. This was done for correct (blue) and incorrect (red) models. The larger the letter, the more significant the bias according to a Fisher's Exact Test.

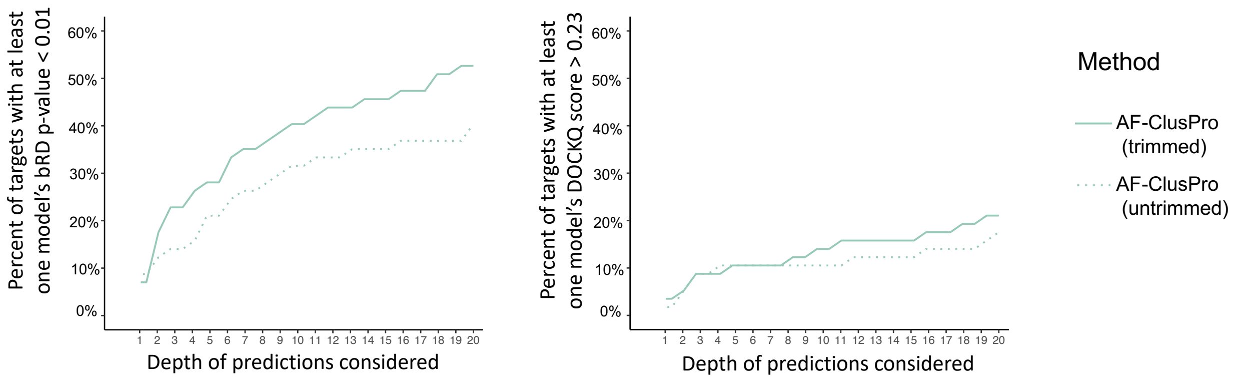


*Figure S8: A Comparison of Running ClusPro with Trimmed or Untrimmed Docking Partners.* Cumulative graphs of the numbers of structures that have at least one correct (dark) or significant (light) structure at various numbers of top predictions are shown. The results of docking with untrimmed antibody and antigen structures are at the top, and of docking with antibody and antigen structures trimmed of residues along each end until a residue of 0.9 or above was hit are at the bottom.

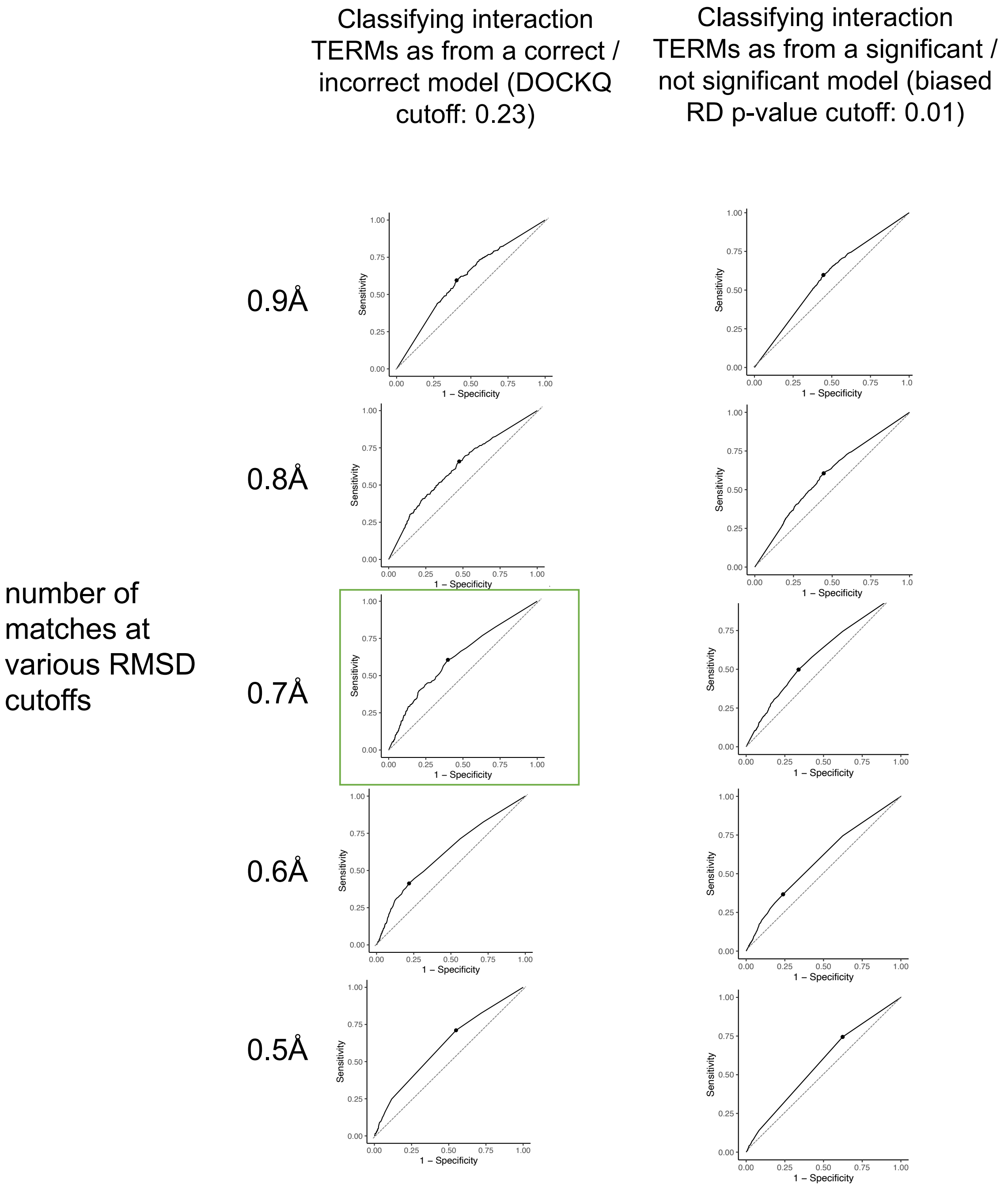


*Figure S9: Determining the Optimal RMSD Cutoff for TERM-based Structural Discrimination*. The interaction-TERMs of all AlphaFold-Multimer top models were searched against a 40% sequence redundancy cutoff database made of the PDB with antibody-like structures removed, to look for the number of matches at various cutoffs. ROC curves to discriminate TERMs as coming from a correct / incorrect structure (left) or significant / not significant structure (right) are shown. The best discrimination according to AUC was 0.7Å used to discriminate correct / incorrect structures (indicated with a green box).

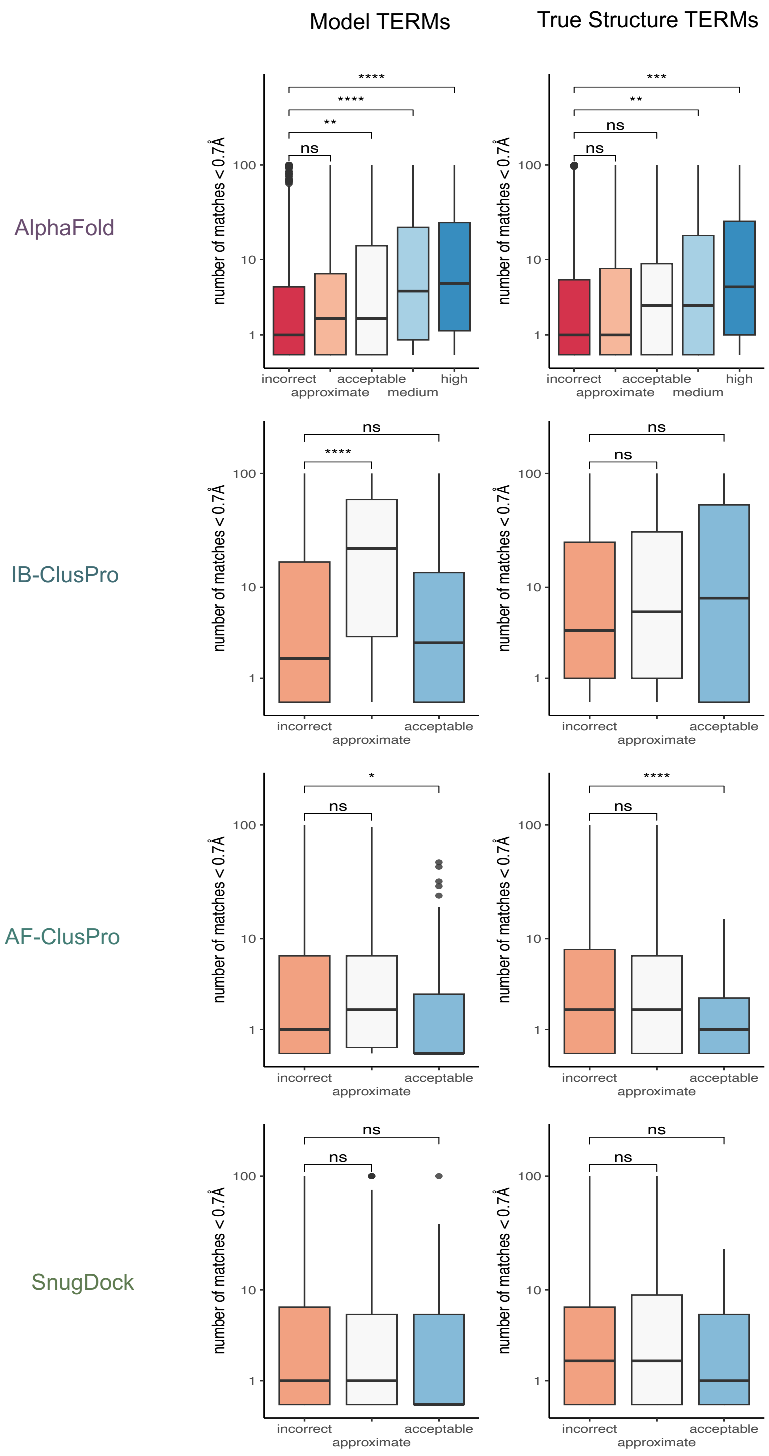


*Figure S10: Structural Biases for Methods Producing At Least One Correct Top Structure, Matches Limited to Interfacial TERMs from the PDB.* Interaction TERMs from each method's models are split depending on if their original model was incorrect, approximate (meaning significant, but not correct), acceptable, medium, or high. Then the number of matches below 0.7*Å* is plotted. This is done for both the TERMs of the model (left) and the TERM of the true structure (right).

Significance between incorrect and other ranks of models are indicated with asterisks (* = < 0.05, ** = < 0.01, *** = < 0.001, **** = < 0.0001).

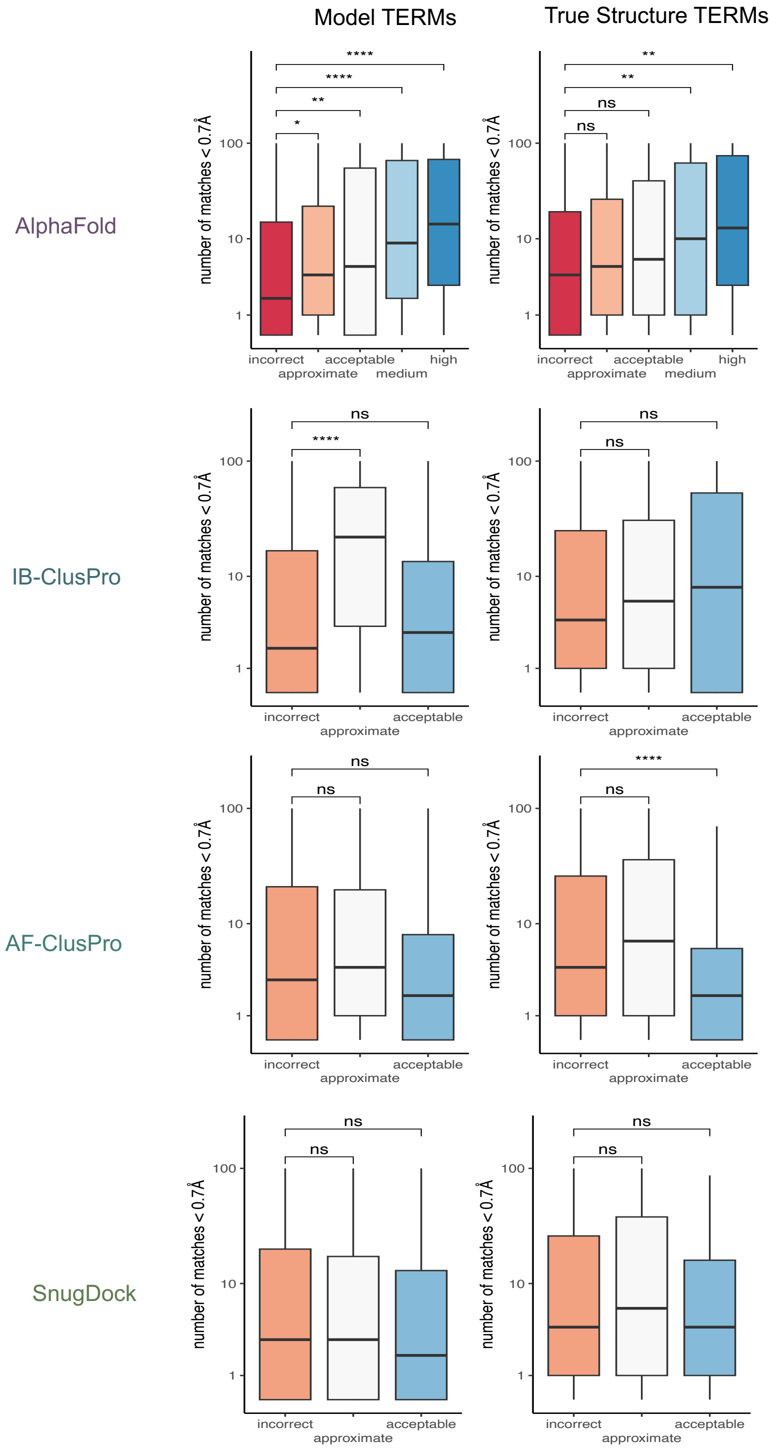


*Figure S11: Structural Biases for Methods Producing At Least One Correct Top Structure, Matches Limited to Non-Interfacial TERMs from the PDB.* Interaction TERMs from each method's models are split depending on if their original model was incorrect, approximate (meaning significant, but not correct), acceptable, medium, or high. Then the number of matches below 0.7*Å* is plotted. This is done for both the TERMs of the model (left) and the TERM of the true structure (right). Significance between incorrect and other ranks of models are indicated with asterisks (* = < 0.05, ** = < 0.01, *** = < 0.001, **** = < 0.0001).
